# Interleukin-1α links peripheral Ca_V_2.2 channel activation to rapid adaptive increases in heat sensitivity in skin

**DOI:** 10.1101/2023.12.17.572072

**Authors:** Anne-Mary N Salib, Meredith J Crane, Sang Hun Lee, Brian J Wainger, Amanda M Jamieson, Diane Lipscombe

**Affiliations:** Department of Neuroscience, Carney Institute for Brain Science, Brown University, Providence, RI 02912, USA; Department of Molecular Microbiology and Immunology, Brown University, Providence, RI 02912, USA; Department of Neurology, Massachusetts General Hospital, Boston, MA 02114, USA

## Abstract

Neurons have the unique capacity to adapt output in response to changes in their environment. Within seconds, sensory nerve endings can become hypersensitive to stimuli in response to potentially damaging events. The underlying behavioral response is well studied, but several of the key signaling molecules that mediate sensory hypersensitivity remain unknown. We previously discovered that peripheral voltage-gated Ca_V_2.2 channels in nerve endings in skin are essential for the rapid, transient increase in sensitivity to heat, but not to mechanical stimuli, that accompanies intradermal capsaicin. Here we report that the cytokine interleukin-1α (IL-1α), an alarmin, is necessary and sufficient to trigger rapid heat and mechanical hypersensitivity in skin. Of 20 cytokines screened, only IL-1α was consistently detected in hind paw interstitial fluid in response to intradermal capsaicin and, similar to behavioral sensitivity to heat, IL-1α levels were also dependent on peripheral Ca_V_2.2 channel activity. Neutralizing IL-1α in skin significantly reduced capsaicin-induced changes in hind paw sensitivity to radiant heat and mechanical stimulation. Intradermal IL-1α enhances behavioral responses to stimuli and, in culture, IL-1α enhances the responsiveness of *Trpv1*-expressing sensory neurons. Together, our data suggest that IL-1α is the key cytokine that underlies rapid and reversible neuroinflammatory responses in skin.

## Introduction

Sensory nerve endings in skin are highly plastic; their sensitivity to stimuli can change rapidly, and reversibly, protecting against potentially damaging insults ^[1]^. Neuroinflammatory responses in skin are triggered by intense or high frequency stimulation of peripheral sensory neurons ^[2]^. A critical rise in intracellular calcium in sensory neurons induces the release of inflammatory signaling molecules, including neuropeptides calcitonin gene-related peptide (CGRP), Substance P, and ATP, all of which have downstream consequences on neuroimmune signaling that underlies the cellular changes associated with inflammation and pain ^[3–7]^. The generation of cytokines from immune cell activation via transmitter receptors such as P2X7, is the key step in defining the inflammatory response including its time course and the involvement of different classes of neighboring sensory nerve endings.

The rapid, adaptive increase in sensitivity to sensory stimuli in skin, in response to potentially damaging events, is one of the best-known examples of neuronal adaptation, but the molecules that mediate the initiating, early steps of the neuroinflammatory response are not fully characterized. In an earlier study, we discovered that voltage-gated CaL2.2 channels expressed in *Trpv1*-nociceptor nerve endings are essential for initiating the rapid neuroinflammatory response to radiant heat following intradermal injection of capsaicin in mice^[8]^.

Ca_V_2.2 channels are abundantly expressed in sensory neurons ^[9–11]^. Our lab, and others have shown that Ca_V_2.2 channels are expressed in sensory neurons of DRG neurons, including in small diameter, capsaicin-responsive neurons, where they contribute to the voltage-gated calcium current ^[8,9,12–21]^. Immunohistochemistry and functional analyses using highly selective conotoxins has shown that Ca_V_2.2 channels are concentrated at a subset of presynaptic terminals in the central and peripheral nervous systems ^[15,22–27]^. In sensory neurons, Ca_V_2.2 channels have long been known to control calcium entry at presynaptic nerve endings of primary nociceptive afferent synapses in the dorsal horn of the spinal cord ^[28–31]^. Intrathecal delivery of highly specific pharmacological inhibitors of Ca_V_2.2 (N-type) channels are analgesic and have been used to treat otherwise intractable pain ^[15,16,32–43]^.Intrathecal ω-CgTx MVIIA rapidly prolonged paw withdrawal latencies to both thermal and mechanical stimuli ^[40]^ consistent with what we observed with global Ca_V_2.2 KO mice (^[8]^ Fig. 2). We previously showed that a local single dose of ω-CgTx MVIIA applied intradermally reduced the robust increase in sensitivity to radiant heat that followed intradermal capsaicin, independent of Ca_V_2.2 channel activity at presynaptic sites in the dorsal horn of the spinal cord ^[8]^ . Interestingly, Ca_V_2.2 channel activity was not essential for the cross-sensitization of mechanoreceptors in the intradermal capsaicin model in the same animals. We also showed that inhibition of P2X7 purinergic receptors, an ATP-gated ion channel that activates immune cells to trigger the release of IL-1 inflammatory cytokines ^[44]^, reduced intradermal capsaicin-induced rapid heat hypersensitivity in skin ^[8]^. The cytokines that link intense activation of *Trpv1*-nociceptors to the rapid, early phase of the inflammatory response in skin, notably sensory hypersensitivity of nerve endings across sensory modalities, are incompletely characterized.

Proinflammatory cytokines such as IL-6, IL-1β and TNFα, are generated during chronic neuroinflammatory responses in skin ^[45–47]^. In addition, IL-1β has been shown to induce changes in excitability of neurons in culture, through effects on ion channel currents and second messenger systems ^[1,48]^. These cytokines are elevated in response to prolonged neuroinflammatory conditions associated with pain, however, most studies measure cytokine levels during the later phases of inflammation, not in the initial acute phase of the rapid hypersensitivity response ^[49,50]^. To address this, we extracted and analyzed interstitial fluid from mouse hind paws within 15 mins following intradermal capsaicin, under conditions that result in rapid behavioral changes in skin to sensory stimuli. Intradermal capsaicin is a well-used model of adaptive sensory hypersensitivity in skin with onset and offset kinetics of tens of minutes. This model parallels short-lasting, heat-induced neuroinflammatory responses in human skin ^[51–53]^.

We show that IL-1L, a proinflammatory alarmin and one of the earliest immune signaling molecules, is generated locally and abundantly in interstitial fluid in response to intradermal capsaicin. The timing of elevated IL-1L levels parallels peak behavioral hypersensitivity to both radiant heat and mechanical stimuli. Our data suggest that IL-1L is both necessary and sufficient to trigger rapid neuroinflammation in hind paws of both *Trpv1-nociceptors* and mechanoreceptors. Our combined findings suggest that co-activation of TRPV1 receptors and voltage-gated CaL2.2 channels are necessary to trigger the release of IL-1L that may subsequently feed back on *Trpv1*-nociceptors to enhance heat responsiveness. Understanding the paracrine signaling, between sensory nerve endings and immune cells in skin during inflammation, will inform strategies to selectively target maladaptive forms of pain while leaving acute protective pain relatively intact.

## Results

Intradermal capsaicin injection induces a robust, rapid, and reversible increase in the sensitivity of the paw to both heat and mechanical stimuli. To identify the cytokines released in the hind paw in response to intradermal capsaicin, and to assess the dependency of cytokine release on global and local Ca_V_2.2 channel activity in skin, we recovered 7-10 µl of interstitial fluid from both paws of each animal, 15 min post injection of capsaicin using three experimental conditions: Intradermal capsaicin in WT, in global Ca_V_2.2^-/-^ KO, and in WT co-injected with Ill-CgTx MVIIA. At the 15 min time point, after capsaicin injection, behavioral responses to radiant heat and mechanical stimuli were enhanced significantly (^[8]^; Fig. 2). The Ca_V_2.2^-/-^ KO mouse strain was generated in our lab and is a global deletion of Ca_V_2.2 (as described previously ^[8]^). Ill-CgTx MIIA is a high affinity, highly selective, and slowly reversible inhibitor of voltage-gated Ca_V_2.2 channels ^[33,54]^. We co-injected Ill-CgTx MIIA together with capsaicin to assess the contributions of peripheral voltage-gated Ca_V_2.2 channels in the skin, locally at the site of injection ^[8,55,56]^.

*IL-1*α *levels are elevated in interstitial fluid in hind paws in response to intradermal capsaicin*

Pilot screens of 20 cytokines were performed on hind paw fluid samples from each experimental condition between two immunoassays, a multiplex bead-based immunoassay (LEGENDplex) and an electrochemiluminescence spot-based immunoassay (MSD, see Methods). Based on these results, and reports in the literature, 13 cytokines were selected as likely neuroinflammatory candidates and screened for using a custom mouse panel LEGENDplex panel. Pooled, paw fluid samples (n = 11-22 mice) were screened. Of the 13 cytokines assayed, only IL-1α was consistently detected in hind paw fluid under any condition at the 15 min time point in both assays. We used a second independent custom panel, Meso Scale Discovery (MSD), to validate IL-1α levels in the same samples (Supplementary Fig. S1). In addition, we assessed: 8 of the same 13 cytokines surveyed on the LEGENDplex, as well as 3 cytokines that were undetectable on our pilot LEGENDplex screen, and macrophage derived chemokine (MDC) which was unique to the MSD platform (Fig. 1a).

**Figure 1.**
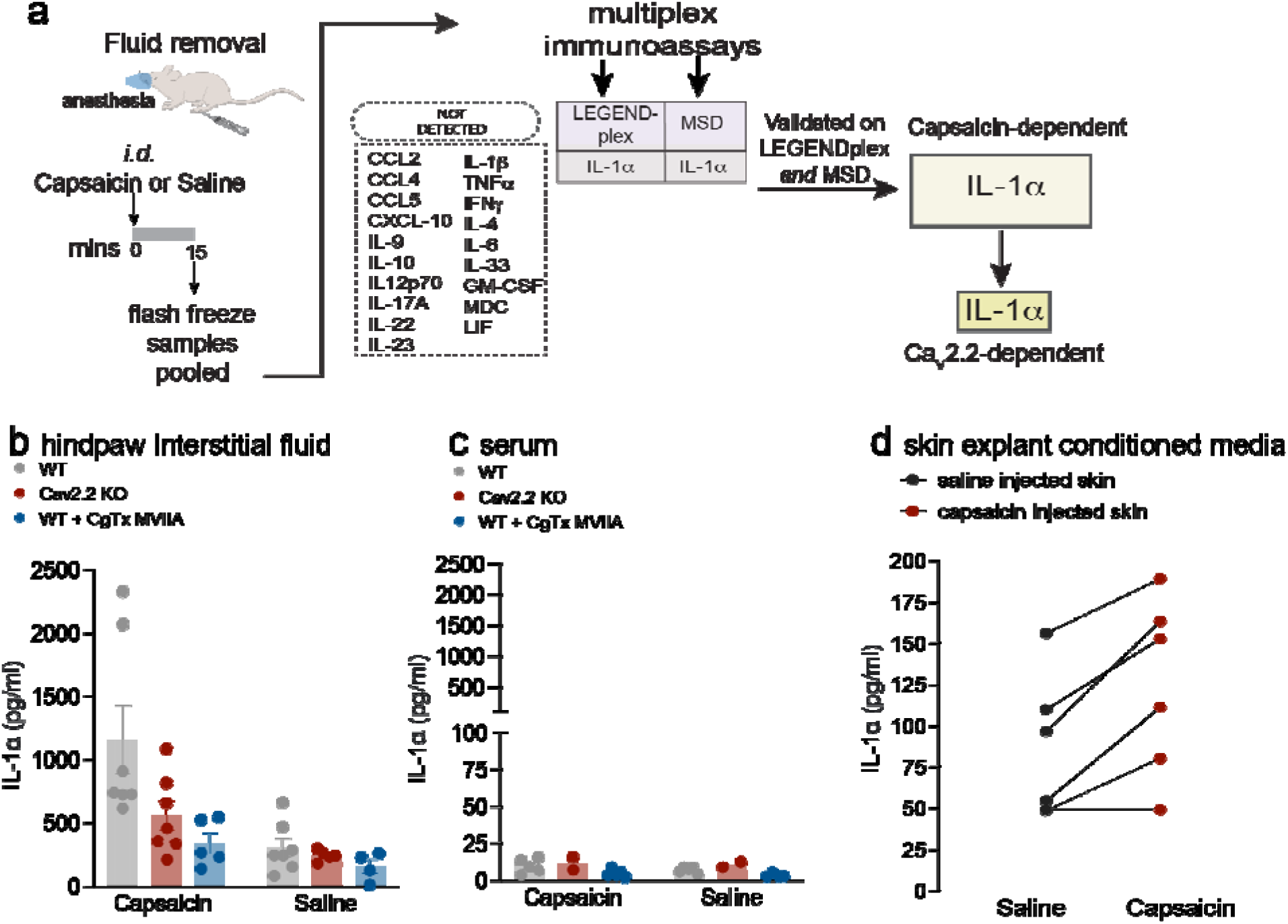
The cytokine IL-1α is present in paw interstitial fluid and its levels increase with intradermal capsaicin and local Ca_V_2.2 channel activity. **a.** Experimental procedures and cytokines screened are shown. Interstitial fluid extracted 15 mins following 50 µl intradermal injection in both paws of 0.1% w/v capsaicin, saline, 0.1% w/v capsaicin with 2 µM ω-conotoxin MVIIA, or saline with 2 µM ω-conotoxin MVIIA. Pooled samples were screened for 20 cytokines listed. **b.** IL-1α levels in hind paw fluid based on panel standards, detected using an inflammatory multiplex bead-based immunoassay (BioLegend, LEGENDplex). Individual points represent pooled samples from wild-type (WT, gray), Ca_V_2.2^-/-^ (KO, red), and WT co-injected with ω-conotoxin MVIIA (CgTx-MVIIA, blue). Number of (pooled samples/mice) for WT with saline (7/18); KO with saline (5/11); WT with capsaicin (7/20); KO with saline (7/17); WT with capsaicin + CgTx-MVIIA (5/14); WT saline with CgTx-MVIIA (4/12). Bars represent the mean for each condition. Mean ± SE for capsaicin: WT = 1159 ± 272 pg/ml; KO = 561 ± 117 pg/ml; WT + CgTx MVIIA = 343 ± 83 pg/ml. p-values calculated by two-way ANOVA with Tukey HSD correction for multiple comparisons hind paw fluid WT capsaicin|WT saline: p=0.0002; KO capsaicin|KO saline: p=0.1600; WT CgTx-MVIIA + capsaicin|WT CgTx-MVIIA + saline: p= 0.4732. WT capsaicin| KO capsaicin: p = 0.0158; WT capsaicin|WT CgTx-MVIIA + capsaicin: p=0.0025;KO capsaicin|WT CgTx-MVIIA + capsaicin: p=0.5875. **c.** IL-1α levels in serum detected using the same cytokine screening protocol as in Fig 1b. Individual points represent pooled samples from the same animals used for hind paw fluid analysis. Bars represent the mean for each condition. Analysis of variance interaction between Injection|Genotype was measured by two-way ANOVA: p=0.8422. **d**. IL-1α levels in skin-explant conditioned media detected using a single analyte ELISA (n=6). Skin punch biopsies were removed 5 mins post-injection and placed in media for 5 mins. Each mouse was injected with saline in one paw and capsaicin in the other for within-animal comparison. Average skin weight for capsaicin injected skin: 5.48 mg; saline injected skin: 4.82 mg. Points represent analysis of conditioned media from each paw and solid lines connect saline and capsaicin injected paws for each mouse; p- value calculated by paired t-test: p= 0.0099.

IL-1α levels were increased significantly in response to intradermal capsaicin in WT mice as compared to saline injected paws (capsaicin vs saline; two-way ANOVA with Tukey correction: p=0.0002; Fig. 1b). The IL-1α response to capsaicin was reduced, but not eliminated, in Ca_V_2.2^-/-^ global KO mice (Fig. 1b; p =0.0158) and also in WT mice co-injected with Ill-CgTx MVIIA (Fig. 1b; p =0.0025), as compared to samples from WT hind paws. We compared the IL-1α levels in hind paws of capsaicin-injected Ca_V_2.2^-/-^ and WT mice co-injected with Ill-CgTx MVIIA mice and these were similar to levels in control hind paws injected with saline (Fig. 1b).

To control for the possible involvement of systemic differences in IL-1α levels across mice under different experimental conditions, we measured IL-1α in serum of the same experimental animals. IL-1α levels in serum were much lower than those at the site of capsaicin injection and they were unaffected by intradermal capsaicin (Fig. 1c). We consistently detected low levels of two cytokines in addition to IL-1α in serum samples, CXCL10 and CCL5, and the levels of these cytokines were not correlated with intradermal capsaicin injection.

In a complementary approach, we measured IL-1α levels using single analyte ELISA levels in hind paw skin-conditioned media. We harvested skin from hind paws of WT mice 5 mins after intradermal capsaicin (ipsilateral) and saline (contralateral) injection. IL-1α levels in hind paw skin-conditioned media from capsaicin injected paws were significantly higher, as compared to those from saline injected hind paws (5 of 6 mice; Fig. 1d; p = 0.01, paired t-test). Collectively, our data show that intradermal capsaicin triggers IL-1α release in hind paws within the first 5 mins of injection, paralleling the rapid time course of behavioral changes in hind paw sensitivities to heat and mechanical stimuli (Fig. 2).

**Figure 2.**
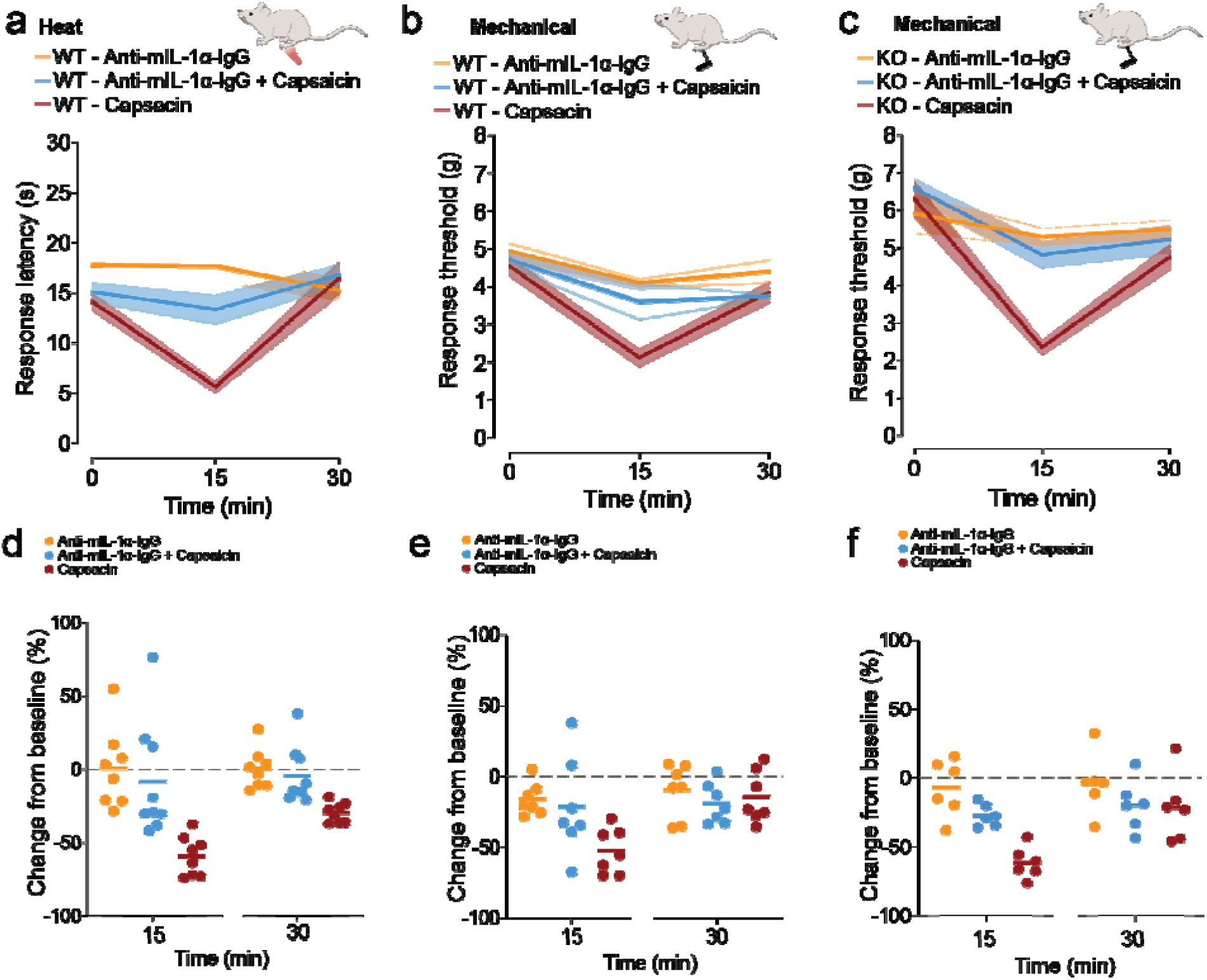
Neutralizing IL-1α in the presence of capsaicin reduces both heat and mechanical hypersensitivity. Paw withdrawal response latencies to radiant heat (**a, d**) and mechanical force (g) using automated Von Frey filament (b-**c, e-f**) represented as mean values (lines) ± SE (shaded area) (*top*; **a,b,c**) and average responses for individual mice (solid circles) and average values for all mice (horizontal line) represented as percent change from baseline (0 mins) (*bottom*; **d,e,f**). Measurements were made immediately prior to (0), and 15 and 30 mins post intradermal (*id*) injections as noted. **a, d.** WT (Ca_V_2.2^+/+^) mice received 20 μL *id* of: 1 μg/ml Anti- mIL-1α-IgG (orange; n=8); 0.1% w/v capsaicin + 1 μg/ml Anti-mIL-1α-IgG (blue; n=9); 0.1% w/v capsaicin (red; n=8). Mean ± SE latencies to heat at 15 mins: 1 μg/ml Anti-mIL-1α-IgG = 17.7 ± 1.6 s; 0.1% w/v capsaicin + 1 μg/ml Anti-mIL-1α-IgG = 13.3 ± 1.5 s; 0.1% w/v capsaicin = 5.6 ± 0.6 s. **d.** 15 min post: capsaicin + 1 μg/ml Anti-mIL-1α-IgG (blue) = -8.35 ± 12.97%; capsaicin = -59.08 ± 4.84%; Anti- mIL-1α-IgG (orange) = 0.88 ± 9.58%. Type | Time interaction p = 0.0008; at 15 min post *id* for: Capsaicin + 1 μg/ml Anti-mIL-1α-IgG | Capsaicin: p=0.0001; Capsaicin + Anti-mIL-1α-IgG | Anti- mIL-1α-IgG p= 0.6887. **b,e.** WT (Ca_V_2.2^+/+^) mice received 20 μL *id* of: 1 μg/ml Anti-mIL-1α-IgG (orange, n = 7); 0.1% w/v capsaicin + 1 μg/ml Anti-mIL-1α-IgG (blue, n = 7); 0.1% w/v capsaicin (red, n = 7). Mean ± SE mechanical response threshold at 15 mins: 1 μg/ml Anti-mIL-1α-IgG = 4.09 ± 0.13 g; 0.1% w/v capsaicin + 1 μg/ml Anti-mIL-1α-IgG = 3.59 ± 0.49 g; 0.1% w/v capsaicin = 2.10 ± 0.24 g. **e.** 15 min post: capsaicin + ml Anti-mIL-1α-IgG (blue) = -21.23 ±13.00%; capsaicin = -52.40 ± 6.04%, 1 μg/ml Anti- mIL-1α-IgG (orange) = -15.90 ± 4.36%. Injection Type| Time interaction: p = 0.0101; 15 min post *id* for: capsaicin + 1 μg/ml Anti-mIL-1α-IgG | Capsaicin: p = 0.0179; capsaicin + Anti-mIL-1α-IgG | Anti- mIL-1α-IgG p = 0.8753. **c,f.** KO (Ca_V_2.2^-/-^) mice received 20 μL *id* of: 1 μg/ml Anti-mIL-1α-IgG (orange, n = 6); 0.1% w/v capsaicin + 1 μg/ml Anti-mIL-1α-IgG (blue, n = 6); 0.1% w/v capsaicin (red, n = 6). Mean ± SE mechanical response threshold at 15 mins: 1 μg/ml Anti-mIL-1α-IgG = 5.29 ± 0.25 g; 0.1% w/v capsaicin + 1 μg/ml Anti-mIL-1α-IgG = 4.82 ± 0.35 g; 0.1% w/v capsaicin = 2.35 ± 0.20 g. Analysis of variance measured by two-way ANOVA and Tukey’s HSD correction for multiple comparisons. **f.** 15 min post: capsaicin + ml Anti-mIL-1α-IgG (blue) = -27.22 ±3.27%; capsaicin = 61.52 ± 4.70%, 1 μg/ml Anti- mIL-1α-IgG (orange) = -7.13 ± 8.38%. Injection Type| Time interaction: p = 0.0175; 15 min post *id* for: capsaicin + 1 μg/ml Anti-mIL-1α-IgG | Capsaicin: p = 0.0081; capsaicin + Anti-mIL-1α-IgG | Anti- mIL-1α-IgG p = 0.1579.

### IL-1α is necessary for the development of capsaicin-induced increases in sensitivity to heat and mechanical stimuli

To establish if IL-1α levels are increased *and* necessary for capsaicin-induced changes in hind paw sensitivity to heat, we co-injected a specific neutralizing monoclonal IL-1α antibody (anti-mIL-1α-IgG, InvivoGen, Catalog # mabg-mil1a) to occlude its actions locally, at the same site of capsaicin injection. We measured hind paw sensitivity to radiant heat and mechanical stimuli in WT mice following capsaicin injection in the absence, and in the presence of anti-mIL-1α-IgG (Fig. 2). Baseline behavioral responses to heat and mechanical stimuli were not consistently affected by anti-mIL-1α-IgG ruling out any direct effect on stimulus-response pathways. By contrast, anti-mIL-1α-IgG reduced significantly capsaicin-induced increases in the sensitivity of mouse hind paws to *both* heat (p = 0.008) and mechanical (p = 0.012) stimuli (Fig. 2a-2e).

In a previous study, we showed that Ca_V_2.2 channel activity was essential for capsaicin-induced hypersensitivity to heat, but notably, not mechanical stimulation, which was fully intact in Ca_V_2.2^-/-^ mice and in hind paws of WT mice injected with ω-CgTx MVIIA. IL-1α levels in response to capsaicin were reduced, but not eliminated, in the Ca_V_2.2^-/-^ global KO mouse (Fig. 1b). We therefore tested the hypothesis that IL-1α levels were still sufficient to mediate capsaicin-induced mechanical hypersensitivity in mice lacking Ca_V_2.2 channels. We used anti-mIL-1α-IgG to neutralize IL-1α in Ca_V_2.2^-/-^ animals and showed that it significantly reduced capsaicin-induced mechanical hypersensitivity (p= 0.0081. Fig. 2c, 2f). The use of anti-mIL-1α-IgG to specifically neutralize the actions IL-1α locally, establish IL-1α as a critical immune signal in the capsaicin-induced neuroinflammatory response in skin.

To establish if IL-1α is sufficient to induce rapid changes in the sensitivity of skin to heat and mechanical stimuli, we directly injected recombinant IL-1α (mIL-1α, R&D catalog #: 400-ML- 005/CF) into the hind paws of mice and assessed their behavior. Paw withdrawal responses to both heat and mechanical stimuli were evoked with shorter latencies (heat) and at lower forces (mechanical) in mice injected with recombinant IL-1α at 15- and 30-min time points post injection, as compared to responses of contralateral paws in the same animals (Fig. 3).

**Figure 3.**
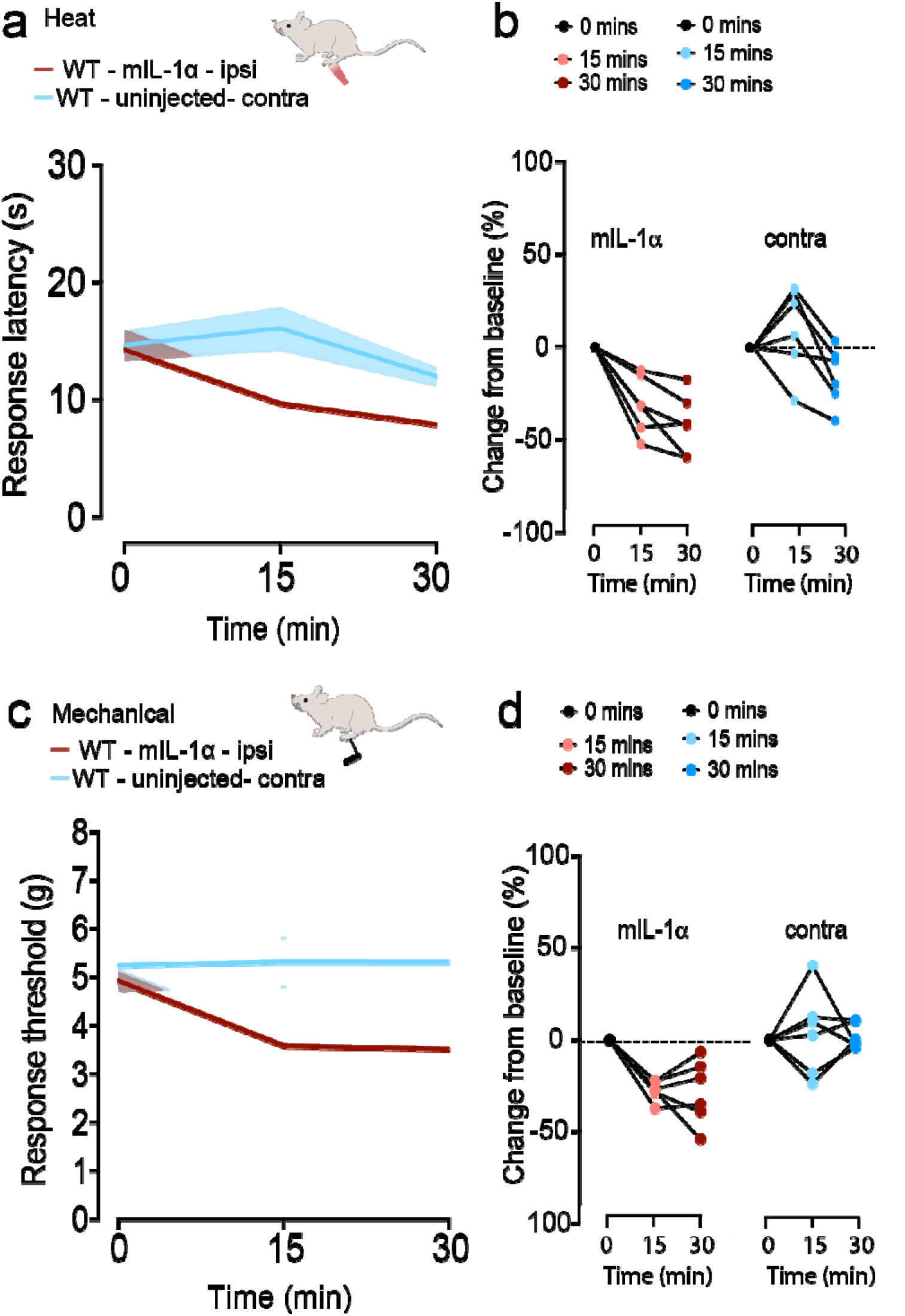
Intradermal recombinant IL-1α triggers both heat and mechanical hypersensitivity in the absence of capsaicin **a**. Response latency (s) to radiant heat measured before (0) and 15 and 30 mins after 20 μg/ml injection of 5 μg/ml recombinant mouse IL-1α (m-IL-1α) in sterile PBS + 10% FBS (n = 6). Injected and un-injected paws (blue). Mean ± SE response latencies to heat 15 mins following injection to: m-IL-1α: 9.66 ± 1.26 s and un-injected 16.06 ± 1.84 s. **b**. Percentage change from baseline for Individual mice in response to m-IL-1a (15 min) -31.03 ± 6.36% and un-injected paws 10.54 ± 10.06 %. Paw injection| time interaction analysis of variance measured by two-way ANOVA and Tukey’s HSD correction for multiple comparisons p = 0.0018. m-IL-1α injected|un-injected p = 0.0075 (15 min) and m-IL-1α injected|un-injected (30 min) p = 0.0235. **c**. Mean ± SE paw withdrawal thresholds to mechanical stimuli before (0) and 15 and 30 mins after 20 μL injection of 5 μg/ml recombinant mouse IL-1α (m-IL-1α) in sterile PBS + 10% FBS (n = 6 mice). Mean ± SE for m-IL-1α: 3.58 ± 0.18 g and un-injected = 5.32 ± 0.52 g. **d.** Average percent change from baseline for each animal at indicated time points. Mean ± SE values at 15 min were m-IL-1a = -27.25 ± 2.22 % and un-injected = 3.78 ± 9.42 %. Paw injection| time interaction analysis of variance measured by two-way ANOVA and Tukey’s HSD correction for multiple comparisons p = 0.0018. At 15 min m-IL-1α injected|un-injected p = 0.0205; 30 min m-IL-1α injected|un-injected p = 0.0068.

Similarly, recombinant IL-1α injected paws were hypersensitive to radiant heat and mechanical stimuli as compared to vehicle injected controls (PBS 10% FBS); Heat: 15 mins mIL-1α compared to vehicle p = 0.07, mechanical at 15 min p = 0.02. Heat: 15 mins mIL-1α compared to vehicle p = 0.07, mechanical at 15 min p = 0.02. Heat: 30 mins mIL-1α compared to vehicle p = 0.005, mechanical at 30 min p = 0.056.

Based on our findings, we conclude that IL-1α is a critical early mediator of transient heat and mechanical hypersensitivity in skin induced by capsaicin. Notably, IL-1α was the only cytokine consistently detected in two different cytokine detection platforms used in our analyses. Our data also show that IL-1α levels are strongly dependent on both TRPV1 receptor activation and peripheral Ca_V_2.2 channel activity, suggesting that IL-1α is a key immune signal underlying the rapid and robust neuroimmune response in skin to heat. Interestingly, even though capsaicin-induced hypersensitivity to mechanical stimulation is independent of Ca_V_2.2 channel activity ^[8]^, we show that this behavioral response also depends on IL-1α.

### Overnight incubation of IL-1α increases responsiveness of sensory neurons

To evaluate the impact of IL-1α on nociceptor responsiveness of primary sensory neurons, we employed APPOINT (Automated Physiological Phenotyping of Individual Neuronal Types), a high-content calcium imaging platform integrating high-throughput single-cell calcium imaging, liquid handling, automated cell segmentation, and analysis including machine learning-based calls of responding cells ^[57]^. Trpv1-Cre mice were crossed with floxed ChR2-EYFP mice and first-generation Trpv1/ChR2-EYFP mice were used for preparation of dorsal root ganglia (DRG) neurons (see methods). We isolated DRG neurons from mice expressing the blue light-sensitive ion channel channel-rhodopsin (ChR2) in *Trpv1*-positive nociceptors and identified optically responsive neurons using a red-shifted calcium indicator. The number and frequency of stimuli (light pulses) in the optogenetic rheobase assay was calibrated to cover the dynamic range of activation ^[57]^. Stimulating TrpV1/ChR2-EYGFP nociceptors with trains comprised of increasing numbers of 1 ms blue light pulses allows us to measure an optical rheobase, the number of stimuli pulses necessary to evoke calcium flux in individual nociceptors. DRG neurons were plated into wells overnight with saline (vehicle) or IL-1α (100 ng/ml). The following day, increasing trains of 1, 5, or 10 pulses of blue light were applied to the wells, and the % of DRG neurons exhibiting suprathreshold levels of calcium flux were quantified. Compared to saline-treated control cells, a larger percentage of cells incubated overnight with IL-1α (100 ng/ml responded to optical stimulation (F_1,40_ = 4.27; *P* = 0.045, two-way ANOVA) (Fig. 4a, 4b). A comparison of the cumulative distribution of Ca^2+^ amplitudes in responsive DRG neurons revealed a slight increase in larger amplitude events in cells treated with IL-1α compared to saline (Fig. 4c), but this difference did not reach statistical significance threshold (*p* = 0.182, two-sample Kolmogorov-Smirnov test). Overall, these data show that IL-1α can act directly on sensory neurons to alter neuronal responsiveness to optical stimulation via the Interleukin-1 receptor type 1 (IL-1R1) receptor on sensory neurons, as has been shown previously for IL-1β^[48]^.

**Figure 4.**
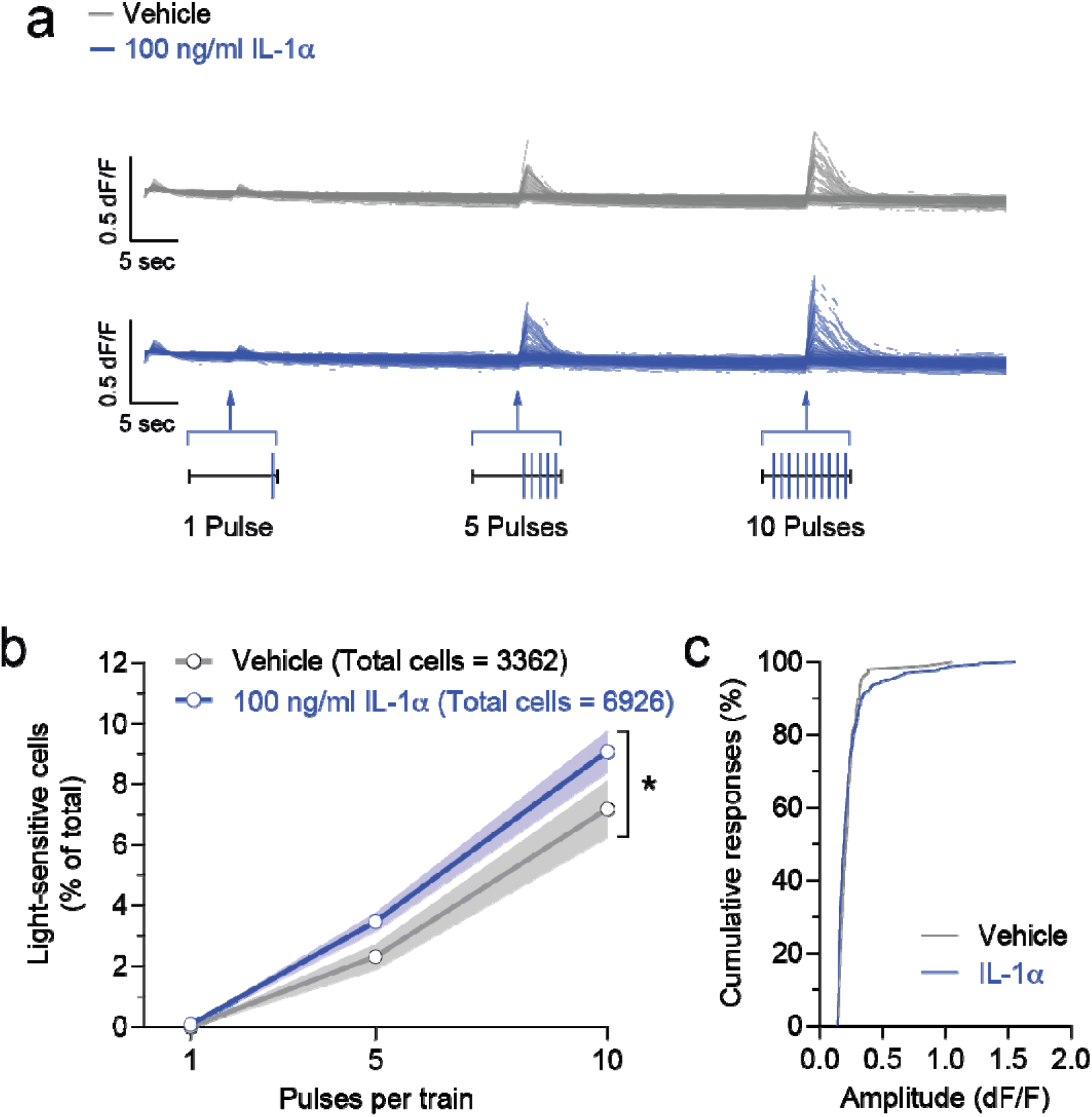
IL-1α reduces the optical rheobase of Trpv1/ChR2-EYFP mouse nociceptors. **a.** Traces of Ca^2+^ responses evoked by blue light stimulations (trains of 1, 5, and 10 1 ms pulses). Scale bars: 5 sec, 0.5 dF/F. **b.** Averaged percentage of sensory neurons activated by increasing pulse number of blue light stimulations following treatment with vehicle and IL-1α (blue: 100 ng/ml IL-1α). Note that overnight incubation with IL-1α significantly increased the percentage of calcium-responsive cells compared to vehicle treatment (F_1,40_ = 4.27; *P* = 0.045, two-way ANOVA with Bonferroni correction for multiple comparisons). Responsive cells were defined as previously using a machine learning-based algorithm ^[57]^ . All data represent mean (point/line) ± SE (shaded area). **c.** Cumulative distributions of calcium response amplitudes measured in DRG somas in response to 100 ng/ml IL-1α (blue) or saline (grey). The Kolmogorov-Smirnov test for comparing two samples did not show a significant statistical difference (*P* = 0.182).

## Discussion

Intradermal capsaicin is used to model one of the most well-known examples of the rapid neuroinflammatory response in skin following exposure to potentially damaging stimuli ^[51,58,59]^. We applied this model in mouse hind paws analyze rapidly generated cytokines in interstitial fluid *in vivo* and, under the same experimental conditions, to monitor behavioral responses to radiant heat and mechanical stimulation. By two independent assays, we identified IL-1α as the major cytokine generated in hind paw fluid and showed it was both necessary and sufficient to support the rapid increase in sensitivity of peripheral nerve endings in skin to sensory stimuli.

We were initially surprised that IL-1α was the only cytokine identified in both cytokine screening platforms and consistently present in mouse hind paw fluid. IL-1α levels were elevated within 15 mins of intradermal capsaicin exposure, coincident with robust behavioral changes to both radiant heat and mechanical stimuli. Levels of IL-1α measured in hind paw interstitial fluid were 3-4 fold higher following capsaicin injection in WT fluid when compared to the dynamic range of IL-1α found in blood in non-injured states ^[60]^. We did not consistently detect other inflammatory cytokines in interstitial fluid samples at the 15 min time point, when behavioral responses are maximal. Importantly, in both assays, we confirmed that IL-6, IL-1β, and TNFα and all other cytokines included in the screen, were detectable in concurrently run standards (Supplementary Fig. S2, Supplementary File S3).

However, our data are consistent with the known role of IL-1α, an alarmin, as one of the earliest signaling molecules of the inflammatory response ^[61]^. In fluid samples of human blisters, IL-1α, not IL-1β, levels are elevated ^[62,63]^. IL-1α is primed and ready in skin resident immune cells for rapid action and, unlike IL-1β which is only biologically active in its cleaved mature form ^[64–66]^, IL-1α activates IL-1R1 in both its cleaved mature and precursor pro forms ^[64]^. The known kinetics of IL-1α are faster than IL-1β following stimulation. Others have shown that on the timescale of hours, IL-1α is maximal within 6 hours of stimulation, while IL-1β is maximal 12-16 hours after stimulation ^[67]^.

We took a number of steps to validate our findings including: i) Employing two independent cytokine detection platforms, Custom Mouse Panel LEGENDplex and MSD U-Plex and R-plex, designed to identify low levels of cytokines in low volume fluid samples; ii) Screening 20 different cytokines; iii) Using single analyte ELISA to measure IL-1α in isolated skin conditioned media; and iv) Using standards to ensure that the cytokine assays are able to detect all cytokines screened, including IL-1β (Supplementary Fig. S2, Supplementary File S3).

To establish if IL-1α is necessary for the rapid behavioral changes in sensitivity to sensory stimuli associated with intradermal capsaicin, we employed a potent neutralizing antibody anti-mIL-1α-IgG. Anti-mIL-1α-IgG is selective for IL-1α, and it inhibits the biological activity of both precursor and mature forms of mouse IL-1α (InvivoGen, Catalog # mabg-mil1a). By contrast, pharmacological inhibition of the IL-1R1 does not distinguish between IL-1α and IL-1β as both cytokines act through the same receptor ^[64,68,69]^. The neutralization of the effects of intradermal capsaicin by anti-mIL-1α-IgG, combined with the demonstration that intradermal recombinant IL- 1α induces heat and mechanical hypersensitivity in mouse hind paw, establish IL-1α as an immune signal that mediates the rapid increase in peripheral nerve ending sensitivity to heat and mechanical stimulation.

In this study, we focus on identifying the cytokine in hind paws that underlies the rapidly developing, and rapidly reversible neuroinflammatory response in skin. Many other cytokines including IL-6, IL-1β, and TNFα are critical for mediating more sustained forms of inflammation associated with chronic pain ^[2,45–47]^. Our data do not exclude the possibility that other cytokines are involved in the capsaicin response in skin, but we do show that IL-1α is a critical signaling molecule in this robust protective response. It is possible that IL-1α has been overlooked as critical for the rapid early phase of neuroinflammation in skin because of a greater focus on chronic inflammatory models of pain, important for identifying new therapeutic targets ^[70–72]^, and the use of assays that do not distinguish IL-1α and IL-1β.

We showed previously that voltage-gated Ca_V_2.2 channels in peripheral nerve endings in skin play a critical role in the development of capsaicin-induced hypersensitivity to heat ^[8]^. Here we show that Ca_V_2.2 channel activity also contributes to increased levels of IL-1α in hind paw fluid in response to intradermal capsaicin. Combining our earlier studies with data presented here suggests a model in which voltage-gated Ca_V_2.2 channels in *Trpv1-nociceptor* nerve endings and IL-1α generating immune cells may function as a signaling unit mediating rapid neuroinflammatory responses that lead to increased heat sensitivity (Fig. 5). Our data lend support to previous studies showing that Ca_V_2.2 channels (N-type) are involved in inflammatory and neuropathic pain responses, as well as in microglial activation and cytokine release ^[73,74]^. It is possible that different classes of voltage gated calcium channels contribute to the generation of IL-1α via mechanoreceptor activation. For example, Ca_V_3.2 channels (T-type), are abundant in low threshold mechanoreceptors (LTMRs) ^[75]^ and they have been implicated in neuroinflammatory pain signaling underlying mechanical allodynia ^[76–78]^. Our conclusions and proposed model (Fig. 5) are the simplest interpretation of data presented here and on published literature: capsaicin acts on TRPV1 receptors expressed by heat responsive sensory nerve endings in skin triggering membrane depolarization, activation of Ca_V_2.2 channels, and release of neuropeptide and ATP that act on immune cells to trigger cytokine release (Fig. 5). But intradermal capsaicin may also act on nonneuronal *Trpv1* expressing cells in skin, including keratinocytes ^[79]^ T-cells ^[80]^ Langerhans cells ^[81]^ which may contribute to Ca_V_2.2-independent sources of cytokines, including IL-1α. For example, capsaicin could act on immune cells directly, triggering the production and/or release of IL-1α independent of Ca_V_2.2 channels and nociceptor activation. We show that IL-1α levels are significantly lower but not eliminated, in hind paw interstitial fluid when Ca_V_2.2 channel activity is inhibited.

**Figure 5.**
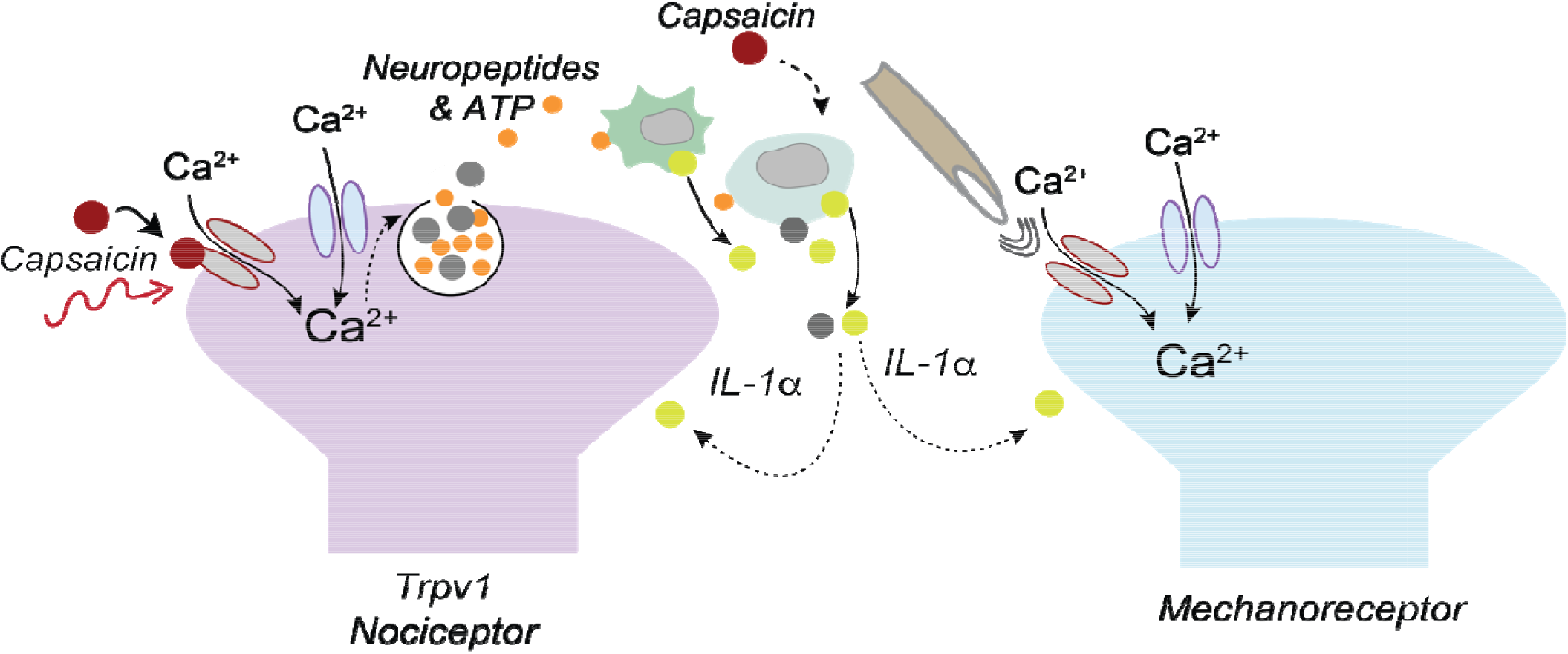
One proposed model for Ca_V_2.2 channel involvement in capsaicin-induced sensory hypersensitivity in skin. This model is based on combined results from published data in DuBreuil et al., 2021 ^[8]^ and this current report. Neuroimmune signaling underlying heat hypersensitivity involves capsaicin activation of TRPV1 receptors on *Trpv1*-expressing cells including nociceptor nerve endings in skin, which induces a membrane depolarization that triggers the opening of voltage-gated Ca_V_2.2 channels. Calcium enters through both Ca_V_2.2 channels and TRPV1 channels, and the influx of calcium through Ca_V_2.2 channels is critical for vesicular release of neurotransmitters. ATP is included based on our previous finding showing that the P2X7 receptor antagonist (A438079) significantly reduces capsaicin-induced heat hypersensitivity ^[8]^, and ATP is known to be present in secretory vesicles ^[10]^. ATP is hypothesized to bind purinergic P2X7 receptors expressed by keratinocytes and leukocytes to trigger release of IL-1α. IL-1α increases the sensitivity of *Trpv1*-expressing nociceptors and mechanoreceptors ^[97]^ to their respective stimuli in glabrous skin. Capsaicin has other targets independent of *Trpv1*-expressing sensory neurons including immune cells, that may contribute to mechanical hypersensitivity which we have shown to be independent of Ca_V_2.2 channel activation. These are indicated as dotted lines as further studies will be needed to dissect the mechanisms underlying capsaicin-induced hypersensitivity to mechanical stimuli.

The cross-sensitization of peripheral nerve endings in skin following intradermal capsaicin or naturalistic stimuli is a classic feature of the rapid protective neuroinflammatory response ^[51,53,82–84]^. Our findings, using the intradermal capsaicin model of rapid inflammation, suggests that IL- 1α also contributes to the recruitment of different sensory modalities through a Ca_V_2.2 channel- independent pathway in addition to *Trpv1*-expressing nociceptors ^[51,85]^. We know from our earlier studies, that capsaicin-induced mechanical hypersensitivity develops independent of peripheral Ca_V_2.2 channel activity in *Trpv1*-nociceptors in the same skin regions ^[8]^. Anti-mIL-1α- IgG occludes both heat and mechanical hypersensitivity induced by capsaicin (Fig. 2a-e), suggesting that sufficient IL-1α must be generated, via a Ca_V_2.2 channel independent pathway, to act on mechanoreceptor nerve endings in skin (Fig. 5). We showed that neutralizing IL-1α in our Ca_V_2.2 global KO mice significantly reduced capsaicin-induced mechanical hypersensitivity (Fig. 2c, 2f) suggesting a concentration dependence of IL-1α for triggering distinct forms of hypersensitivity. Expression levels of IL-1R1 are higher in LTMRs and proprioceptors as compared to *Trpv1*-expressing neurons ^[75]^ therefore, it is also possible that *Trpv1*-nociceptors and LTMRs have different sensitivities to IL-1α.

Others have shown that IL-1R1 receptor activation by IL-1β increases the excitability of cultured neurons isolated from DRG, including via actions on voltage-gated sodium ion channels and potassium channels ^[48,77,86–88]^. As IL-1α acts through the same IL-1R1 receptor as IL-1β, IL-1α should have similar biological effects ^[61,64,66,68]^). We confirmed this using calcium imaging and optogenetic stimulation of *Trpv1*-nociceptors. In this platform we show that overnight incubation of IL-1α increases the responsiveness of cultured *Trpv1*-nociceptors compared to saline-treated control cells (Fig. 4b.) to optical stimulation consistent with other reports using similar methods to assess the actions of proinflammatory cytokines on sensory neurons *ex vivo* ^[89]^. The Wainger lab has previously shown that an increase in the percentage of light sensitive cells as a function of stimulation by blue light reflects increased excitability, as has been observed with potassium channel blockers. Whereas a decrease in the percentage of light sensitive cells as a function of stimulation by blue light reflects decreased excitability, as shown using voltage-gated sodium and calcium channel blockers ^[57]^. Thus, the percentage of neurons responding to optical stimulation by blue light is increased within this sensitive dynamic range by IL-1α. Prior studies have implicated phosphorylation, trafficking, and synthesis of new TRPV1 channels in similar sensitization processes by inflammatory mediators ^[90]^, but further studies are needed to better understand the mechanism of how IL-1a increases neuronal responsiveness. Additional studies are also needed to compare the dose-dependent effects of IL-1α on *Trpv1*-nociceptors and mechanoreceptors that innervate hind paws ^[85,91–93]^.

In conclusion, we report that IL-1α is the critical cytokine released from immune cells in skin, underlying rapid, transient, and adaptive neuroinflammation in skin induced by intradermal capsaicin. IL-1α is necessary and sufficient to couple intense stimulation of *Trpv1* nociceptors to a rapidly developing, but transient, hypersensitivity of local nerve endings in skin to heat and mechanical stimuli. IL-1α participates in cross-sensitization of neighboring mechanoreceptor nerve endings. The *in vivo* actions and time course of IL-1α levels in skin, parallel the behavioral responses that define rapid, adaptive neuroinflammation. Our studies provide important insight for the development of more precise therapeutic strategies to target specific phases of the neuroinflammatory response.

## Materials and Methods

All mice used were bred at Brown University, and all protocols and procedures were approved by the Brown University Institutional Animal Care and Use Committee (IACUC). All experiments were performed in accordance with approved IACUC protocols and compliance with ARRIVE guidelines. Male and female mice were included in all experiments and were 3-6 months old, unless otherwise specified. Values shown are mean ± SE. Experimenters were blind to animal genotype, experimental condition, and solution injected and were only unblinded post analysis, including analysis of interstitial fluid. The Ca_V_2.2^-/-^ global deletion (KO) mouse strain (*Cacna1b*^tm5.1DiLi^, MGI) was generated in our lab by STOP cassette in frame, in exon 1 of *Cacna1b,* as described previously ^[8]^. A wild-type strain was bred in parallel from the same genetic background and used for comparison in some experiments. These wild-type mice were not significantly different in behavioral studies from wild-type littermates and were pooled for analysis. For optogenetic activation of DRG neurons, Trpv1-Cre (Jax 017769) ^[94]^ male mice were crossed with Ai32(RCL-ChR2(H134R)/EYFP) (Jax 024109) ^[95]^ female mice and first-generation Trpv1/ChR2-EYFP pups were used for preparation of primary sensory neurons with ChR2-EYFP expressed in Trpv1-positive nociceptors.

### Hind paw fluid extraction

Mice were anesthetized using isoflurane (2.0-3.5%) and O₂ (0.6-0.8 LPM) administered continuously via nosecone for the entire fluid extraction process. The plantar surface of the footpad was injected with a 30-gauge insulin needle intradermally in the center of the paw with 50 µl of: 0.1% w/v capsaicin, 0.1% w/v capsaicin + 2 µM ω-conotoxin MVIIA, or 2 µM ω- conotoxin MVIIA using saline as vehicle in all solutions. For fluid extraction, the same syringe is used to optimize yield. Fluid is collected by slowly drawing back the syringe to pull any free fluid in the subcutaneous pocket. Light pressure is applied to the surrounding area and any leaked fluid on the surface was collected. Typical fluid yield is 7-10 µl from two paws per animal and samples from 2-4 animals for each condition are pooled into an Eppendorf on dry ice and stored at -80°C until used for immunoassay analyses.

### Serum isolation

Mice were euthanized with an overdose of isoflurane and cervical dislocation and blood was collected postmortem from the heart via cardiac puncture, samples were pooled from the same animals as their corresponding hind paw fluid samples, and after a 45 min coagulation step, centrifuged at 4°C for 12 min at 4000 RPM. Serum supernatant was stored at -80°C for subsequent immunoassays.

### Immunoassays

#### Multiplex bead-based immunoassay (LEGENDplex) fluid analyses

Custom Mouse Inflammation Panel LEGENDplex (BioLegend, LEGENDplex™) protocol was followed according to the manufacturer’s recommendations, and data was acquired using an Attune NxT Flow Cytometer. Pilot hind paw fluid samples from all experimental conditions were collected and used to determine sample dilution factors and to screen for 19 cytokines using capture beads targeting: GM-CSF, IL-1α, IL-1β, IL-4, IL-6, IL-9, IL-10, IL-12p70, IL-17A, IL-22, IL-23, IL-33, TNF-α, IFN-γ, CCL2, CXCL10, CCL4, CCL5, and LIF. We consistently detected only IL-1α in hind paw fluid in the capsaicin model. After initial screening, we selected 13 cytokines for ongoing analyses based on consistent detection in pilot studies and published literature indicating their potential role in neuroinflammation in skin. Biolegend LEGENDplex Data Analysis Software Suite (Qognit) was used to determine analyte mean fluorescence intensities and calculate concentrations based on concurrently-run standard curves (Supplementary File S3).

#### Electrochemiluminescence spot-based immunoassay fluid validation

Following LEGENDplex analyses, remaining samples were assayed on two custom Meso Scale Discovery biomarker panels (MSD R-plex: IL-1α, IL-6; U-plex: TNFα, IL-1β, CCL2, CCL4, CXCL-10, IFNγ, IL-33, IL-10, Il-23, MDC). All samples underwent the same freeze-thaw frequency and duration cycles. MSD Biomarker Group 1 (Mouse) protocol was followed according to the manufacturer’s recommendations for 96 well plate assays. Plates were read and analyzed using an MESO QuickPlex SQ 120MM instrument. Final concentrations reported were adjusted for sample dilution. The concentration of a particular cytokine was determined using a calibrator standard assayed on each plate (Supplementary File S3).

#### Skin explant conditioned media preparation

In anesthetized mice (n=6), we injected capsaicin in one paw, and saline in the contralateral paw prior to skin removal. Mice were euthanized using an overdose of isoflurane followed by cervical dislocation. 5 mins post injection and skin was removed from hind paws using 3.5 mm punch biopsy tools (MedBlade, 2 punches/paw). Skin from saline injected-contralateral and capsaicin injected-ipsilateral paws were incubated separately in 600 µl of prewarmed 37°C RPMI 1610 culture media in 12 well plates. Samples were agitated for 5 min at 250 RPM on an orbital shaker, then 200 µl of conditioned media was removed for analysis. Conditioned media was run in duplicate using an IL-1α ELISA (ELISA MAX Deluxe Set Mouse IL-1α, BioLegend Catalog # 433404).

#### Behavioral assessments

Radiant heat responses were assessed using Hargreaves (Plantar Analgesia Meter IITC Life Science). Mice were placed in Plexiglas boxes on an elevated glass plate and allowed to habituate for 30 min prior to testing. A radiant heat source was positioned beneath the mice and aimed using low-intensity visible light to the plantar surface of the hind paw. For all trials, laser settings were: Idle intensity at 5% and active intensity at 50% of maximum. Cut off time = 30 s. Trials began once the high-intensity light source was activated and ended once the mouse withdrew, shook, and/or licked their hind paw following stimulation. Immediately upon meeting response criteria, the high-intensity light-source was turned off. The response latency was measured to the nearest 0.01 s for each trial using the built-in timer corresponding to the duration of the high-intensity beam. Three trials were conducted on each hind paw for each mouse, with at least 1 minute rest between trials ^[96]^. The average of 3 trials was used for the analysis. N values reported are the number of mice. After baseline measures, mice were anesthetized with isoflurane during all intradermal injections.

Mechanical responses were elicited by an automated Von Frey Plantar Aesthesiometer (catalog #37550, Ugo Basile). Mice were placed in an elevated Plexiglas box with a wire mesh bottom and were allowed to habituate for 30 min prior to testing. The plantar surface of hind paws was assessed using a steady ramp of force ranging from 0 to 8g for up to 90 sec. The trial is automatically terminated when the filament buckles or the paw is withdrawn, force and reaction time are captured. After baseline measures, mice were anesthetized with isoflurane during all intradermal injections.

#### Primary DRG harvesting and culture

For optogenetic calcium imaging experiments, male and female mice (C57Bl6/J) between 14-21 days old were used. *Trpv1*/*ChR2-EYFP* strains were generated by crossing *Trpv1-Cre* (Jax 017769) male mice with LSL-*ChR2-EYFP* (Jax 024109) female mice. C1-L6 dorsal root ganglia (DRG) were dissected from isolated and bisected spinal columns of postmortem mice. DRG were placed in cold DMEM/F-12 media (Thermo Fisher 11320033) and transferred to collagenase/dispase for 60 minutes at 37°C. To dissociate cells, a series of mechanical trituration steps were performed. Cells were filtered through a strainer, centrifuged using a BSA gradient to separate out debris, and plated on a poly-D-lysine (PDL)/laminin treated 96-well plate. All primary sensory neurons were imaged after being cultured overnight in neurobasal media (Thermo Fisher 211 03049) supplemented with B27 (Thermo Fisher 17504044), Glutamax (Thermo Fisher 35050061), and Penicillin-Streptomycin (Thermo Fisher 15070063).

#### Calcium imaging assay

Calcium imaging experiments were performed using an ImageXPress micro confocal high content imaging system (Molecular Devices). Cells were incubated with CalBryte-630AM (AAT Bioquest 20720, 3 µg/mL in 0.3% DMSO) calcium indicator for 30 minutes in the dark. Media was then replaced with 100 µL physiological saline (in mM: 140 NaCl, 5 KCl, 2 CaCl_2_, 1 MgCl_2_, 10 D-glucose, 10 HEPES, and pH 7.3-7.4 with NaOH) immediately prior to cell imaging. For optogenetics experiments, three trains of blue light pulses (1, 5, and 10 pulses per train; 1 ms each at 50 Hz) were delivered and cells were imaged using the APPOINT calcium imaging assay, as described previously ^[57]^.

### Data quantification and statistical analysis

Initial identification and quantification of CalBryte-630AM intensity was performed using a custom journal in ImageXPress (Molecular Devices) analysis software. Briefly, CalBryte-positive cells in each well were identified using a minimum projection of the timelapse image stack based on size and CalBryte intensity. Each cell automatically generated an individual ROI, which was transferred back to the original timelapse stack and the mean intensity of each ROI was calculated for every image. Analysis of cell intensity and quantification was performed in R (Version 4.3.1). Statistical analyses were performed using RStudio (Posit Software), Prism (Version 10; GraphPad), or Excel (Microsoft). All data are presented as the mean ± SE. The significance of optical rheobase assay was assessed using two-way ANOVA followed up by Bonferroni’s multiple comparisons.

## Supporting information

Supplementary Figure S1 and Annotation legend for Supplementary File S2

## Data Availability Statement

The Ca_V_2.2^-/-^ mouse strain (Cacna1b^tm5.1DiLi^) is described in the MGI database and available by request to Diane Lipscombe. All datasets are available by request to Diane Lipscombe.

## Competing interests

The author(s) declare no competing interests that might be perceived to influence the results or discussion reported.

## Author contributions

A-M.N.S, M.J.C, S.H.L, B.J.W, A.M.J and D.L. contributed to research design; A-M.N.S, M.J.C, and S.H.L performed research experiments; A-M.N.S, M.J.C, S.H.L, and D.L. analyzed data; A-M.N.S and D.L wrote the manuscript. All authors reviewed the manuscript.

## Funding

This work was supported by NINDS NS055251 (D.L.); NHLBI R01HL165259 (A.M.J.), NHLBI R01HL126887 (A.M.J.), P20GM121344 Pilot Project (A.M.J.), Carney Innovation Award (A.M.J.); NIA 5R21AG075419 (B.J.W.)

## Notes

### Competing Interest Statement

The authors have declared no competing interest.

### Summary of Updates

Figure 2 revised to include new data; Funding information added; Supplemental files updated.

