## Supplementary Figure S1 and Annotation legend for Supplementary File S2 for "Interleukin-1α links peripheral Ca_V_2.2 channel activation to rapid adaptive increases in heat sensitivity in skin"

**Figure S1**

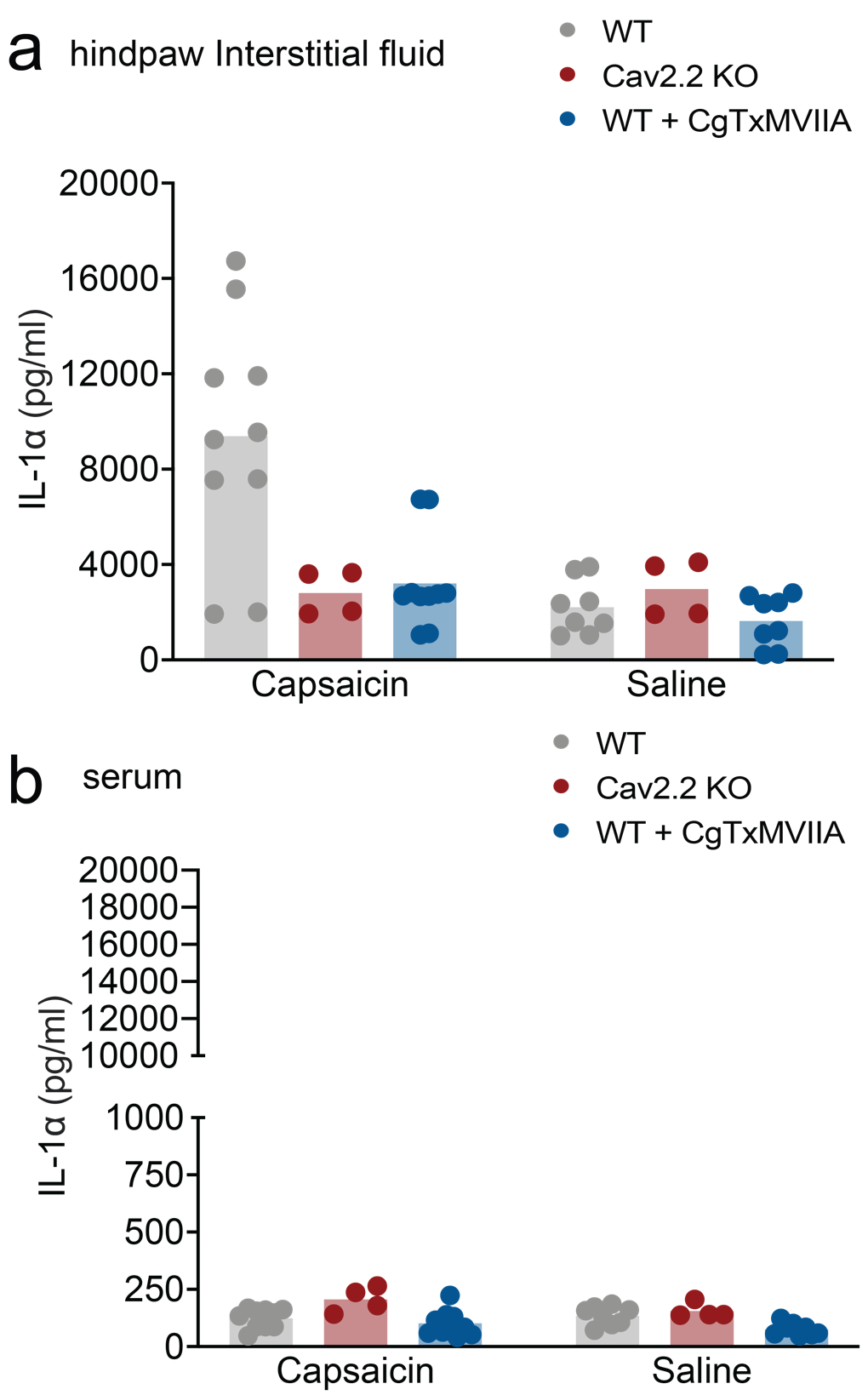

**Figure S1:** Meso Scale Discovery (MSD R-plex and MSD U-Plex) electrochemiluminescent immunoassay. **a-b**. Validation of relative IL-1α levels measured using the MSD immunoassay in hind paw fluid (a) and in serum (b) from the same fluid extracted from hind paws as detailed in Fig. 1b. and Fig. 1c. in the main body of the report. Data shown are from two technical replicates.

**Figure S2**

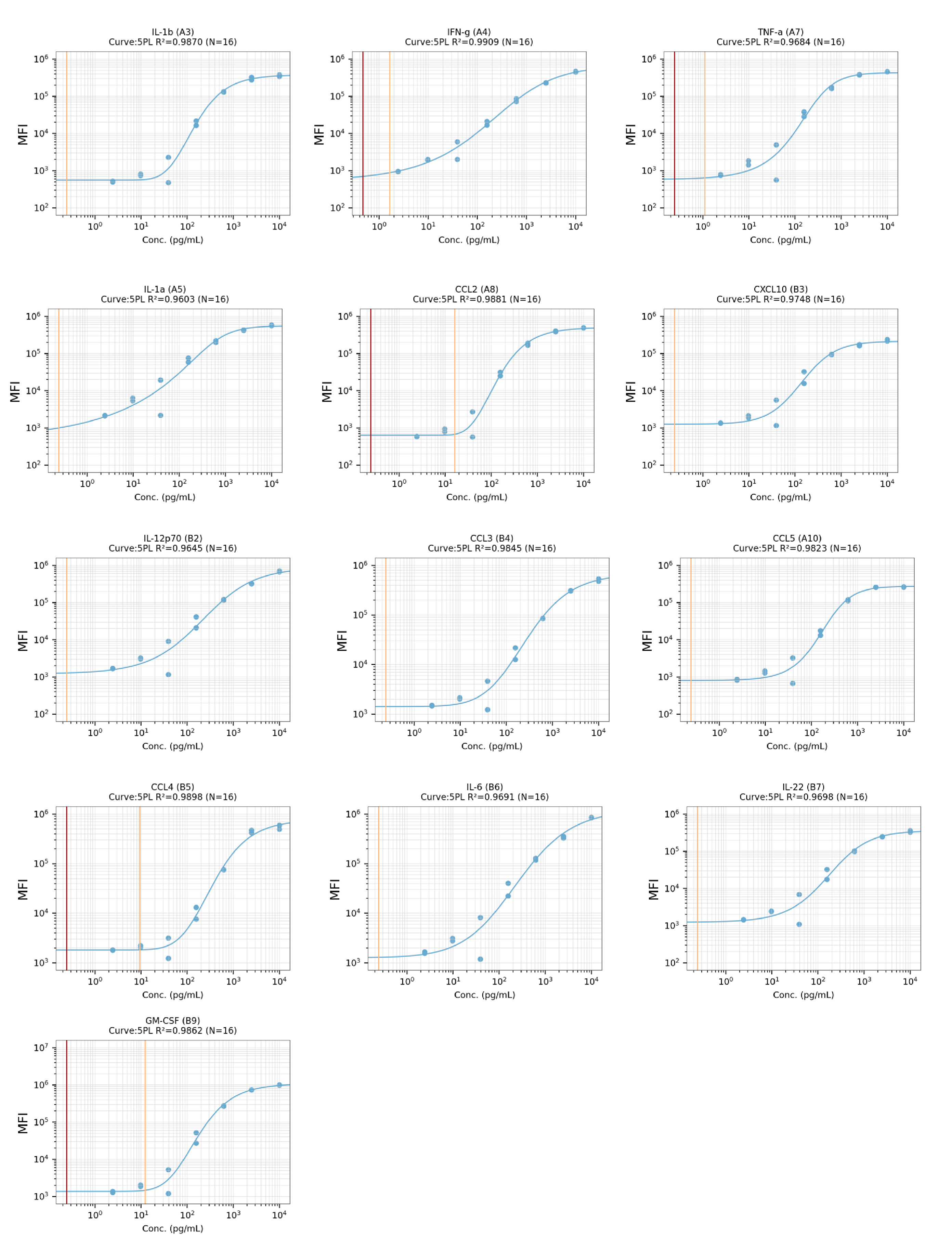

**Figure S2** Concurrently run standard curves generated for each of the 13 analytes assessed in using a custom multiplex bead-based immunoassay (LEGENDplex, see methods). Two technical replicates for each of the 8 standards were measured and fit to a 5PL curve. Standards serve as a positive control for analyte detection and are used to determine the detection range.

**Annotation for Supplementary File S3**

**Output from Biolegend LEGENDplex Qognit software and MSD Quickplex software.** IL-1α was the only cytokine consistently detectable in hind paw fluid samples across both platforms**.**

**LEGENDplex Summary:** Capture bead IDs corresponding to distinct beads of difference sizes used to identify each analyte, IC50 values for each analyte, and R^2^ values generated for concurrently run standards.

**LEGENDplex-output**: Predicted concentration using mean fluorescence intensity signal reported in picogram/ milliliter (pg/ml) for each of the analytes measured based on concurrently run standards. Sample types include standards, hind paw fluid, and serum.

**MSD-output**: Calculated concentration reported in picogram/ milliliter (pg/ml) for each analyte spot using electrochemiluminescence signal and whether the sample was the detection range of the assay based on concurrently run calibrator standards. Sample types include standards, hind paw fluid, and serum.
